# MassDash: A Web-based Dashboard for Data-Independent Acquisition Mass Spectrometry Visualization

**DOI:** 10.1101/2024.01.15.575772

**Authors:** Justin C. Sing, Joshua Charkow, Mohammed AlHigaylan, Ira Horecka, Leon Xu, Hannes L. Röst

**Author notes:** Equal Contribution (Justin C. Sing, Joshua Charkow).

## Abstract

With the increased usage, diversity of methods and instruments being applied to analyze Data-Independent Acquisition (DIA) data, visualization is becoming increasingly important to validate automated software results. Here we present MassDash, a cross-platform, DIA mass spectrometry visualization and validation software for comparing features and results across popular tools. MassDash provides a web-based interface and Python package for interactive feature visualizations and summary report plots across multiple automated DIA feature detection tools including OpenSwath, DIA-NN, and dreamDIA. Furthermore, MassDash processes peptides on the fly, enabling interactive visualization of peptides across dozens of runs simultaneously on a personal computer. MassDash supports various multidimensional visualizations across retention time, ion mobility, m/z, and intensity providing additional insights into the data. The modular framework is easily extendable enabling rapid algorithm development of novel peak picker techniques, such as deep learning based approaches and refinement of existing tools. MassDash is open-source under a BSD 3-Clause license and freely available at https://github.com/Roestlab/massdash, and a demo version can be accessed at https://massdash.streamlit.app.

## Introduction

Data-Independent Acquisition (DIA) is a robust mass spectrometry (MS) technique that provides a thorough analysis of complex biological samples. DIA systematically samples fragment ions leading to high coverage and quantification accuracy,^1,2^ making DIA a valuable technique for a wide variety of applications^3–5^. The proteomics field is widely adopting DIA for its ability to provide in-depth and accurate insights about complex biological systems with applications ranging from cancer research to systems biology^3,6–9^. Its adoption is also met with continuous advancements in mass spectrometry technology, experimental design and sophisticated data analysis algorithms^2,10–14^. For example, the original peptide-centric DIA analysis strategy^2^ continues to be refined with novel methods of library generation,^12^ deep learning based peak pickers,^15^ and statistical scoring^13,16,17^. Furthermore, library-free approaches, using deep learning predicted peptide libraries^13^ or spectrum-centric approaches^18,19^ have also increased in popularity. Additionally, these approaches can be applied to novel instrumentation and sampling strategies^20–27^ and chemical multiplexing^28,29^. Considering the diverse strategies and applications in which DIA is being applied, there is likely not one best analysis workflow for every application or every instrument. Given the complexity of DIA data, it is unlikely that there is a universal set of parameters within a tool that can be applied ubiquitously. Moreover, the abundance of available tools may overwhelm individuals, potentially steering them toward suboptimal workflows. Thus, it is more important than ever for researchers to visualize the raw data as quality control to ensure that software functions as expected for the given context.

Although several visualization tools for DIA experiments exist, each has its own set of limitations. TOPPView^30^ and Skyline^31^ are well-established open-source visualization and data exploration tools with vibrant communities. However, to prioritize performance, these tools are developed in low-level languages that make it challenging for novice users to prototype new features for novel data analysis. TAPIR^32^ and AlphaViz^33^ are two visualization tools written in Python, a user-friendly high-level language, however, they come with limitations including the requirement to store data locally in the case of TAPIR or limited support to an individual commercial vendor form in the case of AlphaViz. Furthermore, both TAPIR and AlphaViz only support OpenSwath and DIA-NN outputs respectively. To that end, we sought to provide a user-friendly, Python-based DIA visualization tool that supports quick visualization from several automated feature detection tools and enables on-the-fly testing and prototyping of new algorithms.

We present MassDash, a novel platform tailored for researchers and analysts in the dynamic realm of DIA mass spectrometry. With a lightweight visualization library, it excels in prototyping algorithms (i.e., peak picking, scoring and multi-run alignment), scaling with multiple runs, and efficiently reading results from common DIA feature detection tools. Seamlessly integrated with Streamlit, MassDash enhances the user experience, offering a powerful graphical user interface (GUI) for algorithm testing and parameter optimization. Accessible both locally and remotely through a web browser, this platform offers versatile data accessibility. Locally, it optimizes performance by leveraging high-speed network connections and minimizing input/output (I/O) latency for swift data processing. Remotely, it excels in managing large datasets, ensuring enhanced accessibility and scalability, which significantly contributes to collaborative research efforts. Furthermore, MassDash also integrates with Jupyter Notebooks to allow for reproducible scripting of visualizations and rapid algorithm development in Python. MassDash’s versatility aligns with modern research needs, emphasizing flexibility and accessibility in data analysis workflows. MassDash provides advanced capabilities for chromatogram visualization, algorithm testing, and parameter optimization, thus empowering researchers to customize analyses and streamline workflows.

## Methods

### Benchmarking Datasets

#### SWATH-MS Gold Standard (SGS) Dataset

In this study, we utilized sixteen DIA-MS runs from the SWATH-MS Gold Standard (SGS) dataset introduced by Röst. et. al. 2014. These runs comprise eight samples containing 0% *S. pyogenes* in human plasma and eight samples containing 10% *S. pyogenes* in human plasma. The SGS dataset encompasses 422 chemically synthesized stable isotop-labeled standard (SIS) peptides.^2^ The DIA-MS runs were acquired on an AB SCIEX TripleTOF 5600, and acquisition parameters are as described in the supplement of Röst. et. al. 2014^2^. Raw mzML DIA data, sqMass files and OpenSwath (OSW) peak boundaries were acquired from Gupta. et. al. 2023^34^. Briefly, raw MS files were converted to the mzML using MSConvert without centroiding. Automated targeted analysis was performed using the OpenSwathWorkflow tool from OpenMS, using optimized parameters as described in the methods of Gupta. et. al. 2023^34^. For identification and quantification comparison, DIA-NN (version 1.8.0) was run on the same 16 runs to identify features using the default settings and the same library used for the OpenSwathWorkflow analysis.

#### Two-Proteome diaPASEF Dataset

To demonstrate MassDash’s visualization capabilities for ion mobility enhanced MS data, six diaPASEF files (Bruker .d) were acquired from a published two-proteome dataset from Meier et. al. 2020^20^. Briefly, the two-proteome dataset consists of peptides from HeLa and yeast cell lysate mixed at different ratios: sample A (HY_200ng_45ng) contains 200ng human and 45ng of yeast protein lysate, and sample B (HY_200ng_15ng) contains 200ng of human and 15ng of yeast protein lysate. The diaPASEF runs were acquired on a Bruker timsTOF Pro, and acquisition parameters are as described in Meier et. al. 2020^20^. The Bruker vendor tdf files were converted to mzML using diaPysef (version 1.0.10). OpenSwath results showcased are from the original publication^20^. DIA-NN analyses were conducted for the sample A triplicates and sample B triplicates. DIA-NN analysis was conducted on version 1.8.0 with the two proteome spectral library in the original publication^20^ published in Meier et al. 2020 via the command line interface with the following settings: --reanalyse, --smart-profiling, --matrices, --report-lib-info --matrix-qvalue 0.05.

#### Synthetic-Phospho DIA Dataset

In phosphopeptide datasets, peptidoforms commonly elute close to one another meaning that standard peak picking parameters might not be appropriate.^35–37^ To showcase the importance of validation and the potential for parameter optimization for peak-picking algorithms, we utilized a single run (highest dilution sample) of a synthetic-phospho peptide dataset from Rosenberger. et. al. 2017^35^. Briefly, 579 synthetic heavy-isotope labeled phosphopeptides were spiked into a human cell line background in a 13-step dilution series^35^. Spectral library generation and OpenSwathWorkflow analysis are as described in the original publication, and the resulting library and results files were acquired from the public PRIDE repository (PXD004573). OpenSwathWorkflow was re-run using the same parameters as in the original publication to generate the extracted ion chromatogram (sqMass) file.

#### Benchmarking Code Execution Time Performance and Memory Usage

The major computation steps: feature identification data access, MS data access initialization, MS data extraction, plot generation and plot rendering were tracked and recorded for 200 random peptide precursor selection interactions of MassDash’s GUI. These major components were measured as time execution blocks, from the beginning of code execution until the end, using python’s built-in module timeit. To measure memory usage, the GNU tool/usr/bin/time was used to measure the maximum amount of resident memory used by MassDash’s GUI for 10 and 200 random interactions with the GUI (random precursor plotting).

## Results and Discussion

### MassDash’s Framework and Purpose

The increasing utilization and technical advancements in high-throughput MS, particularly in DIA, have driven the development of novel algorithms for data processing and deconvolution. However, the plethora of tools and workflows presents a problem for researchers attempting to identify the optimal approach for their data. We therefore present MassDash, a software platform for visualization and algorithm development capable of cross-workflow visualization and validation of DIA results with a capable Python back-end library for rapid iteration of algorithm development to support novel workflows (Fig. 1). MassDash uses raw MS data files and output from commonly used automated DIA analysis software for real-time, interactive visualization of peptide precursor signals. Additionally, MassDash also plots summary statistics including coefficient of variation distributions and identification rates. This comprehensive and flexible platform addresses critical needs in the field, offering visualization and validation capabilities for existing tools and serving as a robust framework for generating algorithms tailored to the challenges posed by high-throughput mass spectrometry data.

**Figure 1.**
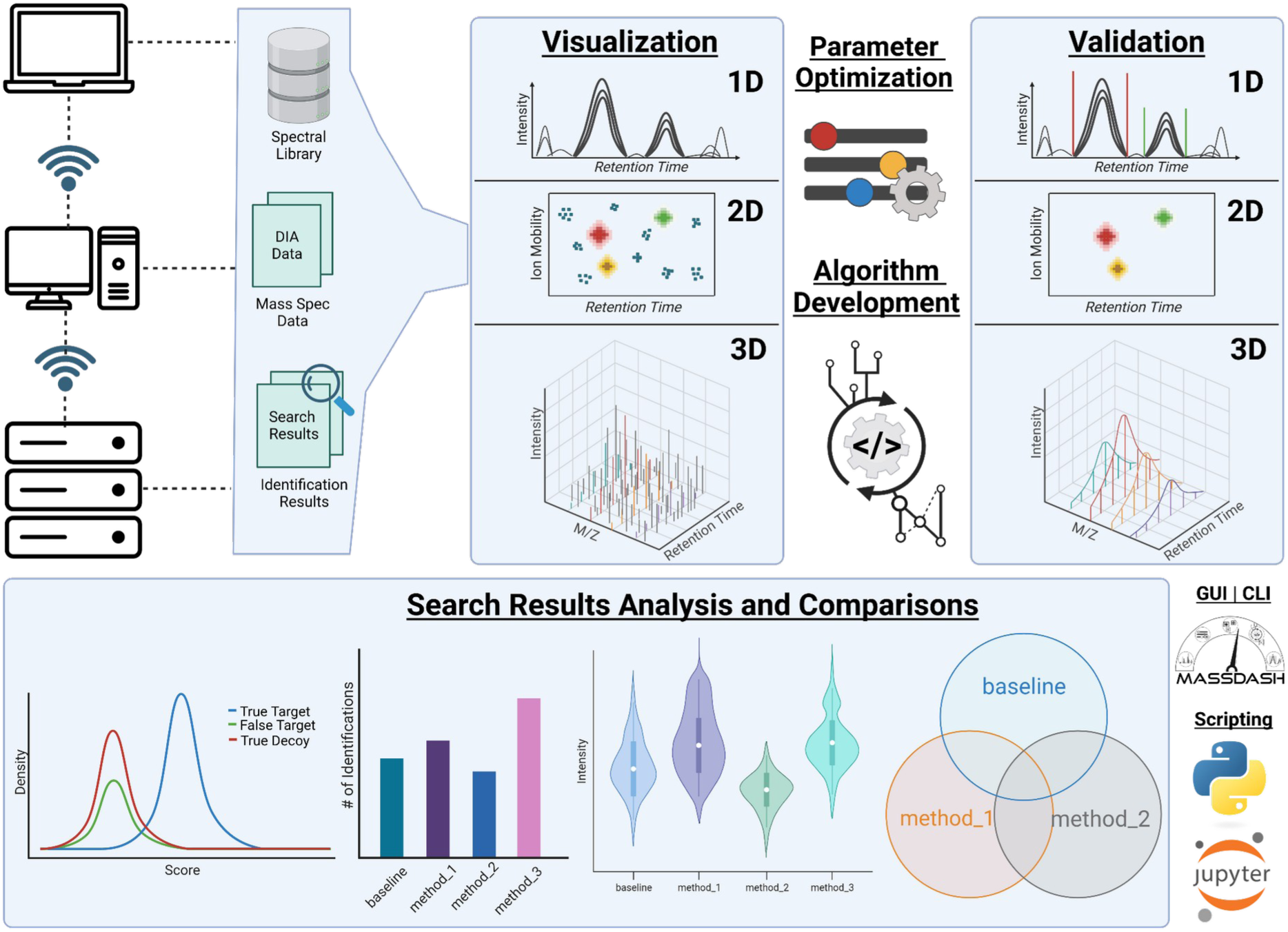
MassDash: A Comprehensive Toolkit for Streamlined DIA-MS Visualization, Analysis, Optimization, and Rapid Prototyping. This schematic provides an overview of the MassDash toolkit and its versatile workflows. Accessible through a graphical user interface (GUI) deployed as a web application either locally or remotely, MassDash requires DIA/diaPASEF MS data, whether raw (.mzML) or in the form of post-extracted ion chromatograms (.sqMass), a spectral library, and output from an automated DIA processing tool (OpenSwath, DIA-NN or dreamDIA). The tool offers diverse visualization options, including 1D, 2D, or 3D plots tailored to the data type. Raw data parameter optimization empowers users to finely tune and explore the dataset before initiating comprehensive targeted data extraction. Summary statistics can also be compared including coefficient of variation, identification rates, and overlap in identifications between software tools or runs. Beyond its user-friendly interface, MassDash serves as a Python library, facilitating rapid algorithm development and testing. Users can delve into and compare search results derived from different methods, enhancing the tool’s utility for robust data exploration and analysis. Created with BioRender.com (2024).

MassDash is a modular framework, allowing easy integration of new functionalities and algorithms. Leveraging the Streamlit library, MassDash provides a user-friendly GUI accessible via a web browser. Written in Python, it seamlessly integrates with other Python packages, enhancing its capacity for rapid algorithm development—particularly valuable for testing and implementing deep learning algorithms. Moreover, MassDash functions not only as a GUI but also as a versatile Python package that can be seamlessly incorporated into scripting environments, including Jupyter Notebooks (Sup. Fig. 5). Furthermore, MassDash utilizes Sqlite and pyOpenMS^38^ for efficient and low-memory data access and retrieval across multiple mass spectrometry runs. The analysis in Supplementary Table 1 shows consistent memory usage across varying file sizes and data types, indicating effective memory management within the application. Memory consumption remains stable around 2 GB for sqMass files, regardless of the number of input files (1 to 16). Similarly, memory usage for DIA mzML and diaPASEF mzML files also increments by a small amount despite the number of input files and interactions. Benchmark tests on major computing operations reveal that data extraction is the most time-consuming process, averaging around 0.71 seconds for a single post-extracted ion chromatogram file (XIC, sqMass) and 7.97 seconds for the 16 runs from the SGS dataset post XIC data (Fig. 2, Sup. Fig. 1, Sup. Table 2). When analyzing and visualizing raw DIA data, the extraction time is approximately 5.26 seconds for a single file and 9.42 seconds for two files. This demonstrates that if you have pre-extracted sqMass files, MassDash can efficiently and quickly extract and render XICs. In contrast, extraction from raw data requires more computation, resulting in a longer wait time; however, it still completes within seconds.

**Figure 2.**
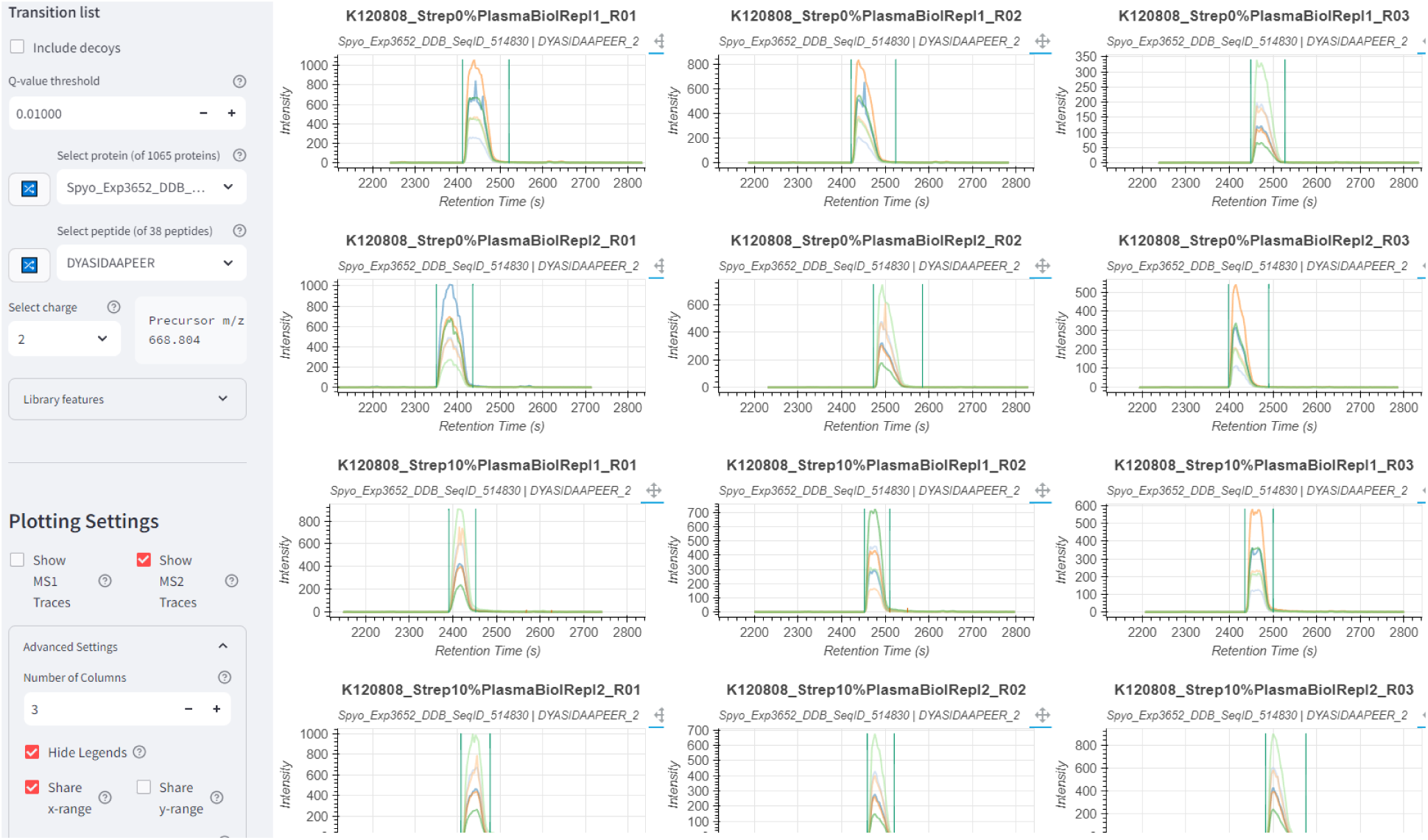
Visualization and Validation of Multiple Experiments with MassDash. Extracted Ion Chromatograms (XICs) are displayed for the protein Elongation factor Tu (peptide: DYASIDAAPEER, charge +2) across 16 runs. The left sidebar allows users to choose the protein/peptide for plotting XICs and select different available peak-picking methods for assessment. While most runs consistently pick the most intense peak, it is noticeable that the peak in the first run is split in half by the peak-picking software (OpenSwath). This highlights the importance of validating software tools involved in data processing to ensure result quality. The data used in this example is sourced from the gold standard *S. pyogenes* data, available on PeptideAtlas (PASS01508).

### The Importance of Validation and Visualization

Many automated feature detection tools typically incorporate methods to control the number of false discoveries, thereby offering a degree of confidence in the resulting identifications.^39^ However, current statistical approaches in the field use a family-wise error rate (FWER) which does not provide a probability for an individual peak to be correctly picked. This works well for global analyses, but is problematic if only a few proteins are selected for follow-up analyses (as in many wet-lab biological experiments or biomarker studies) since the false positive rate among a selected subgroup of data points could be much higher than among the total number of peptides^40^. Therefore, manual validation of results is crucial to ensure that quantifications obtained from data processing tools accurately extract signals for the correct peak. This involves confirming that peaks are not incorrectly cut off, avoiding the return of questionable peaks, and ensuring consistent selection of the same peak across multiple runs (Fig. 2). This is evident in Figure 3, which depicts instances where automated software tools align with manual inspection (Fig. 3a), situations where tools converge on the same feature, but manual validation reveals it as questionable due to a strong precursor signal but has weak fragment ion signals (Fig. 3b), and scenarios where tools identify entirely different features (Fig. 3c). MassDash facilitates the validation of feature detection tools by overlaying their detected features onto the extracted data. This capability extends beyond a single dimension, enabling the visualization of two to three-dimensional plots to assess the performance of data extraction parameters across different dimensions, including mass-to-charge, retention time, and ion mobility (Sup. Fig. 3, Sup. Fig. 4).

**Figure 3:**
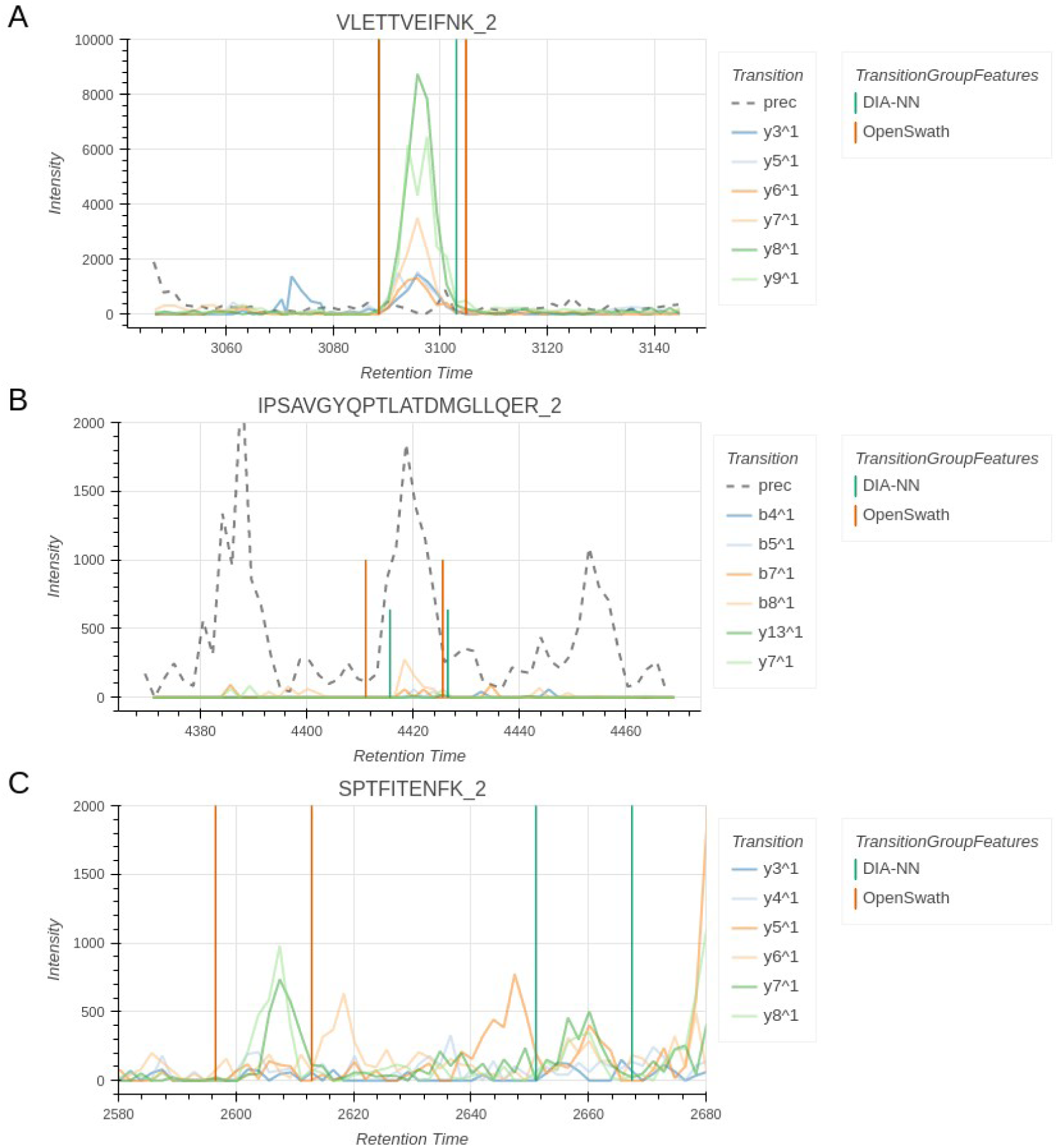
Automated feature detection tools are not always straightforward. Chromatograms of select precursors in a replicate of sample B from the two-proteome diaPASEF dataset with precursor boundaries shown for DIA-NN (green) and OpenSwath (orange). Titles above each chromatogram plot indicates the peptide and the charge state. A) Example of a precursor where automated software tools are in agreement with manual inspection. B) Example of a precursor where both automated feature detection tools significantly identify a questionable precursor. Although this precursor has a strong precursor signal, the fragment ion signals are weak and provide questionable evidence of the precursor’s presence. C) Example where automated feature detection tools identify different features for the same precursor. Both features chosen by the software tools look reasonable for this noisy precursor. Manual visualization is helpful to determine which feature, if any, to accept as the true feature.

It is crucial not only to visually validate the detected features but also to compare different feature detection tools because this comparative analysis enables assessment of an algorithm’s performance, data specific parameter optimization, tool selection, evaluate reproducibility, and address algorithmic biases (Fig. 3, Fig. 4, Sup. Fig. 2). Given the multiplexed nature of DIA data and the continuous development of novel tools, it becomes essential to compare and evaluate different methods. In our benchmarking SGS dataset, we illustrate the comparison between three feature detection tools—OpenSwath, DIA-NN, and DreamDIA—at a 1% FDR at the peptide level (Sup. Fig 2). Across 16 runs, both deep-learning algorithms (DIA-NN, DreamDIA) identify roughly the same number of 11,000 peptides, while OpenSwath identifies around 9,800 peptides. Notably, OpenSwath and DreamDIA exhibit higher variance in the number of identifications across the 16 runs, whereas DIA-NN shows a much lower variance. Regarding the coefficient of variation (CV), their performance is fairly similar, although DIA-NN and DreamDIA have a greater number of outliers with a CV above 250. Additionally, the comparison reveals peptides uniquely identified by each software tool, emphasizing the importance of visual validation to explore intriguing case examples and understand the nuances of identification methods.

**Figure 4:**
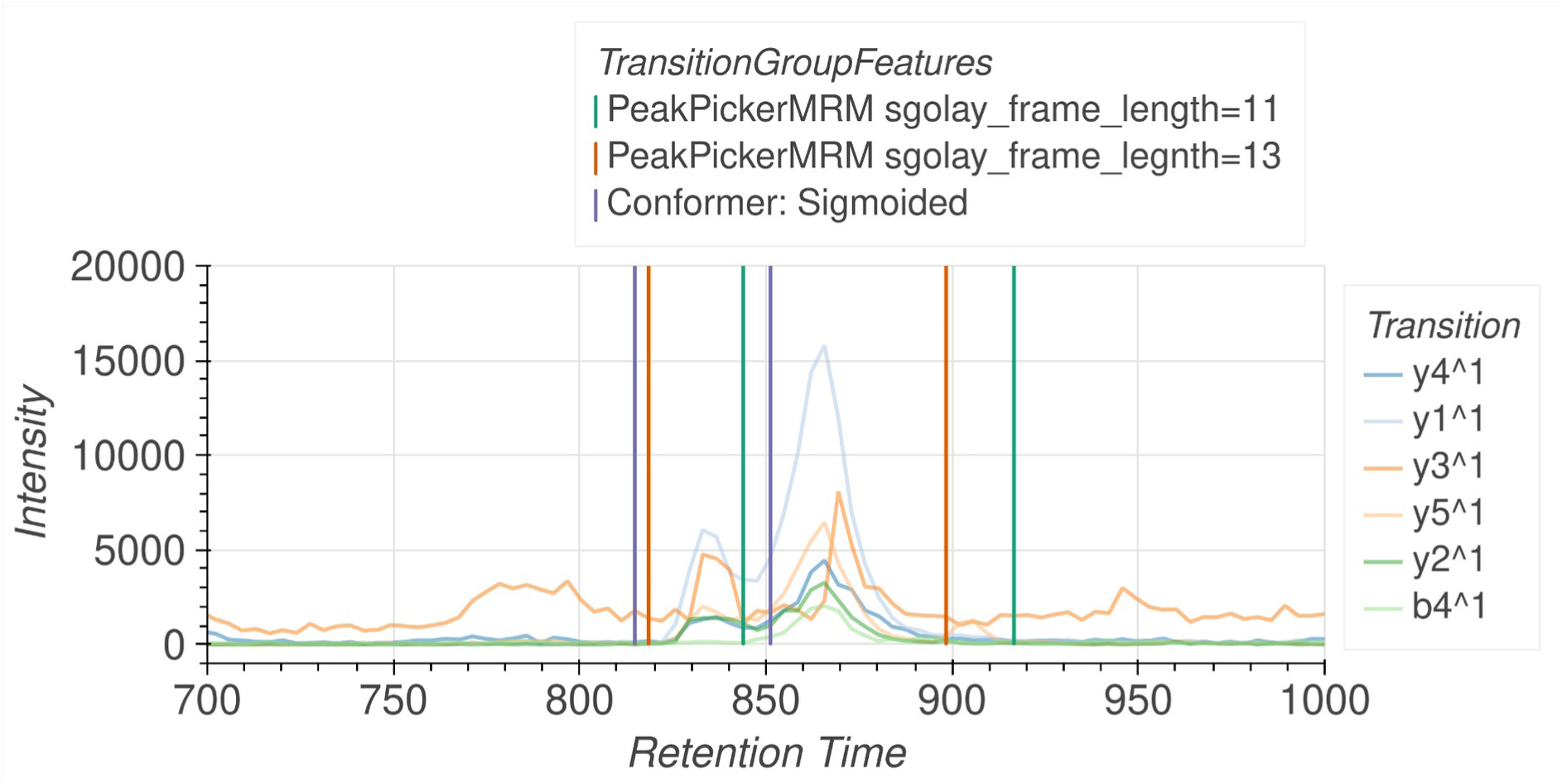
Algorithm Development With MassDash. Chromatogram of peptidoforms *NKESPT(Phospho)KAIVR, charge +3* and *NKES(Phospho)PTKAIVR, charge +3*. In this chromatogram, these two peptides are closely eluting with one another. Three different peak pickers are shown the MRMPeakPicker (OpenSwath default peak picker) with an sgolay_frame_length of 11, MRMPeakPicker with a sgolay_frame_length of 15 and the deep learning Conformer peak picker. Each peak picker picks different peak boundaries, with the PeakPickerMRM sgolay_frame_length=13 grouping the two peptides into one peak, indicating the need to optimize peak pickers for specific experimental contexts.

### MassDash as a tool for Parameter Optimization

With the diverse application of DIA to many different contexts and instruments, the default settings for a given software tool may not always be appropriate for the given experiment. MassDash’s plug-and-play functionality provides a conducive environment for the swift integration of different algorithms, streamlining the process of exploring and optimizing analytical approaches. This flexibility enables researchers to effortlessly implement and test various algorithms for diverse tasks, including feature extraction, peak picking, and other DIA-MS analysis methods of interest.

For example, the optimal extraction parameters across retention time, m/z, and ion mobility may be different for different experiments or even for different precursors in the same experiment. Although enlarging extraction windows leads to higher sensitivity, this comes at the cost of a decrease in selectivity. Since MassDash contains built-in peak pickers and can perform on-the-fly extraction, different extraction parameters can be tested and visualized in an interactive environment. For example, in OpenSwath, peak picking is performed by extracting and summing up signals across a specific area of ion mobility followed by peak picking on the resulting chromatogram. A commonly used ion mobility extraction window for diaPASEF data is 0.06 1/k0 since this provides buffer room to account for differences between the library and feature ion mobilities. However, in a sample A replicate from the two-proteome diaPASEF, for precursor *NAFLESELDEKENLLESVQR, charge +3* the extraction window of 0.06 appears too large since there is interference in the precursor signal leading to an overestimation in the peak boundaries (Fig. 5A). Visualizing the 2D heatmap, it is clear that there is an interfering precursor above (*NPSFSEEPWLEIK, charge +2)* and that refining the ion mobility extraction window should improve the chromatogram signal-to-noise (Fig. 5B, 5C). After refining the ion mobility extraction window the transition chromatograms are cleaner allowing for a more accurate quantification estimation of peak boundaries (Fig. 5D).

**Figure 5:**
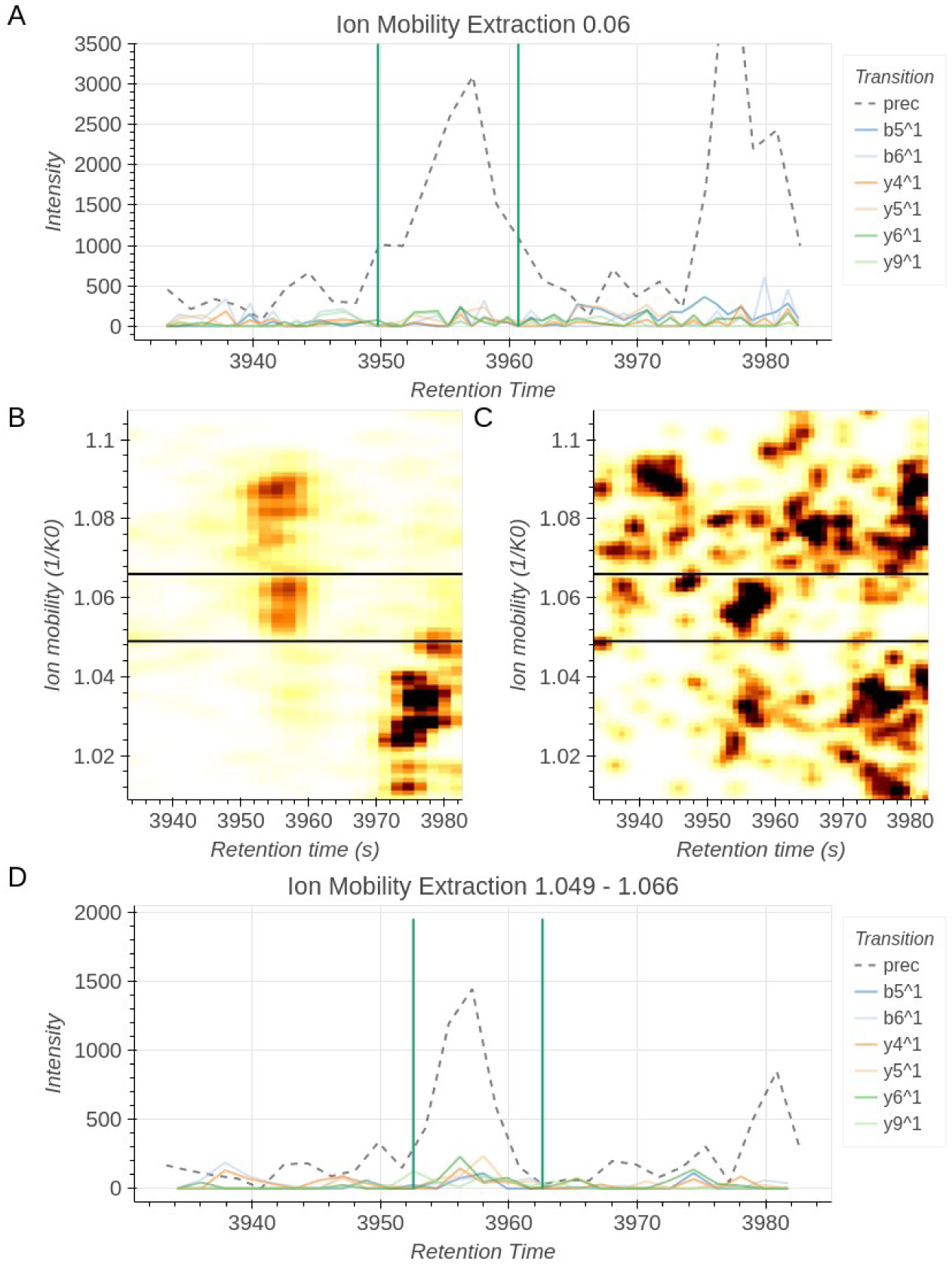
Multidimensional plotting can be used for extraction width optimization. A) Chromatogram of precursor *NAFLESELDEKENLLESVQR, charge +3* with an ion mobility extraction width of 0.06, centered around the precursor (1.028 - 1.088). This is the extraction window commonly used for OpenSwath to account for variations in ion mobility. The fragment ion signals are very noisy and the precursor signal contains an interfering peak on the left side leading to an overestimation of the peak group width. B) 2D chromatogram (retention time vs ion mobility) with Gaussian blur and histogram equalization of the MS1 signal of precursor *NAFLESELDEKENLLESVQR, charge +3*. Through visualization, the extraction window (black box) was manually drawn to exclude the precursor above (*NPSFSEEPWLEIK, charge +2*). C) 2D chromatogram with Gaussian blur and histogram equalization of the amalgamated MS2 transition signals. Ion mobility boundaries used for MS1 are in agreement with MS2 2D chromatogram. D) Chromatogram of precursor *NAFLESELDEKENLLESVQR, charge +3* with an ion mobility extraction of 1.049 - 1.066. Boundaries were manually determined from figure B) and C). With the updated extraction window the precursor and fragment ion peaks are cleaner leading to easier detection of the peak group boundaries.

Additionally, MassDash’s Python backend facilitates seamless integration with numerous Python packages enabling rapid method prototyping and testing. For example, MashDash supports the native OpenSwath peak picker meaning that peak picker parameter optimization can be done interactively and the new optimal parameters can be used in a subsequent OpenSwath analysis. One instance in which peak-picking parameters may need to be adjusted is in phosphoproteomic datasets. These datasets have a high prevalence of peptide isoforms which due to their similarity may partially coelute. This partial co-elution would require less smoothing to be performed by the peak picker to obtain accurate boundaries. For example, using the OpenSwath peak picker with high Savitzky–Golay smoothing, the peak picker may mistakenly group the two coeluting peptides into one peak (Fig. 4, orange line). Decreasing the smoothing parameter allows the peak picker to correctly distinguish between the two partially coeluting peaks (Fig. 4, green line).

### Novel algorithm development

MassDash’s modularity and Python interface promote the rapid development of novel tools, including deep learning-based peak pickers. This empowers researchers to test and validate machine learning and deep learning algorithms effortlessly within a ready-to-use modular environment. Many AI frameworks, being in Python, further facilitate integration with MassDash, enhancing accessibility and streamlining the incorporation of cutting-edge methodologies. MassDash’s adaptability extends to the utilization of saved trained models, providing a valuable tool for researchers to assess the performance of machine learning and deep learning algorithms on their specific datasets. This feature enhances the platform’s utility in exploring cutting-edge approaches, advancing algorithm development, and contributing to the ongoing evolution of DIA-MS methodologies. To demonstrate this, we have built-in support for the Conformer deep learning peak picker^15^. Interestingly, when applying the Conformer peak picker to the phosphopeptide chromatogram above, the small peak is chosen as the peak of highest confidence in contrast to the OpenSwath peak picker (Fig. 4, purple line). This might be because the Conformer peak picker also takes into account library retention time and relative ratios of fragment ions, whereas OpenSwath’s peak picker prioritizes intensity. MassDash’s object-oriented codebase allows users to easily extend the application and implement and test their own peak picker.

## Conclusions

In conclusion, we present MassDash, a novel modular software framework for mass spectrometry DIA visualization, which offers a streamlined, user-friendly interface for rapid quality control visualization, prototyping, and algorithm testing. Its Python library integrates seamlessly into scripting environments like Jupyter Notebooks, enhancing accessibility and adaptability. MassDash’s capacity for on-the-fly data processing, efficient memory usage and, rapid algorithm development in Python distinguishes it from existing tools, providing researchers with a versatile platform for visualization and validation of DIA data. Overall, MassDash’s practical features make it a valuable asset for researchers navigating the complexities of high-throughput mass spectrometry data analysis.

## Data Availability

Initial implementation and testing was performed using the gold standard *S. pyogenes* dataset, available on PeptideAtlas (PASS01508), as well as a synthetic phosphorylation dataset available on PRIDE (PXD004573). Testing of raw targeted extraction for diaPASEF data was done using a two-proteome mixture dataset available on PRIDE (PXD017703). Ion mobility test files were created from the 50ng dilution set (PXD017703, 20190816_TIMS05_MA_FlMe_diaPASEF_25pc_50ng_A2_1_26.d)

## Acknowledgments

We are thankful to Röst lab members for testing, validation and input throughout the development of the tool. We would also like to thank Mingxuan Gao for his input and help with incorporating DreamDIA into the MassDash. J.C.S. was supported by the European Research Area Network Personalized Medicine Cofund (PerProGlio). J.C. was supported by CIHR (#506078) and the Ontario Graduate Scholarship. M.A. was supported by the Chan Zuckerberg Initiative (#513040). I.H. was supported by Early Research Award (ERA) (#511828) and the University of Toronto Open Fellowship. L.X. was supported by NSERC (#506685).

## Contributions

J.C.S., J.C., and H.L.R. designed the study. J.C.S. and J.C. designed and developed the software and M.A, L.X. and I.H. contributed to development. J.C.S., J.C., M.A, L.X., I.H. and H.L.R. contributed to writing and analysis. H.L.R. supervised the study.

