## Supplemental Text for "MassDash: A Web-based Dashboard for Data-Independent Acquisition Mass Spectrometry Visualization"

### I. Supplemental Discussion

#### A. Justification for Creating a New Visualization Software

Several tools are available for chromatogram visualization in the realm of DIA experiments. Skyline, for instance, is a well-established tool with a vibrant community, primarily used for chromatogram visualization in DIA MRM and PRM experiments[1]. However, its rich feature set may be daunting for new users, making it difficult for new users to rapidly prototype programmatically and it has limitations such as being supported only on Windows and requiring all data to be stored locally. TAPIR provides a lightweight solution as a Python-Qt-based software for visualizing extracted ion chromatograms and scored peak groups in DIA analysis[2]. It includes an executable that can run on the three major platforms (Windows, Mac OS, and Linux). However, it's important to note that the data must be stored locally, and the tool lacks the capability to allow on-the-fly testing of parameter tuning for targeted extraction analyses or rapid prototyping of data processing algorithms. TOPPView, as part of the OpenMS suite, serves as a visualization tool for mass spectrometry data, offering a user-friendly interface for data exploration;[3] however, its functionality may be limited to the capabilities within the OpenMS library. TOPPView was initially designed for small-scale SRM/MRM applications and may face challenges with high-throughput data[2], limiting its suitability for rapid prototyping or on-the-fly parameter tuning. Finally, there is AlphaViz, an open-source Python package that facilitates the visualization and validation of protein and peptide identifications in ion mobility-enhanced DIA-MS data[4]. While providing a chromatogram view and real-time ion mobility visualization, it is limited to supporting the Bruker TimsTOF proprietary vendor (.d) data format. Additionally, it has memory efficiency limitations, as it loads the entire dataset into memory, which may prevent users from viewing multiple files simultaneously for cross-run comparison.

#### II. Supplemental Method Notes

##### Internal Subsampled Unit Test MS Data Files

To perform continuous integration and deployment unit testing, as well as to provide small example demo datasets, MassDash provides small test files created from publicly available datasets. Benchmarking sqMass files of a synthetic phosphopeptide dataset were acquired from the PRIDE archive (PXD004573) and were subsampled for a few precursor examples using SQLite and pandas in Python. Ion mobility test files were prepared using the published 50ng HeLa cell dilution dataset from Meier et al 2020 (*20190816_TIMS05_MA_FlMe_diaPASEF_25pc_50ng_A2_1_26)* and its corresponding published and newly processed results. The OpenSwath (.osw) file was created by converting and subsetting the published .tsv results file (pyprophet_export_25pc_50ng.tsv). The DIA-NN report file was created by processing using the settings outlined above. Test .mzML files were created by converting the Bruker .d file using diaPysef (version 1.0.10) and filtered using pyOpenMS.

**III. Supplemental Figures and Tables**

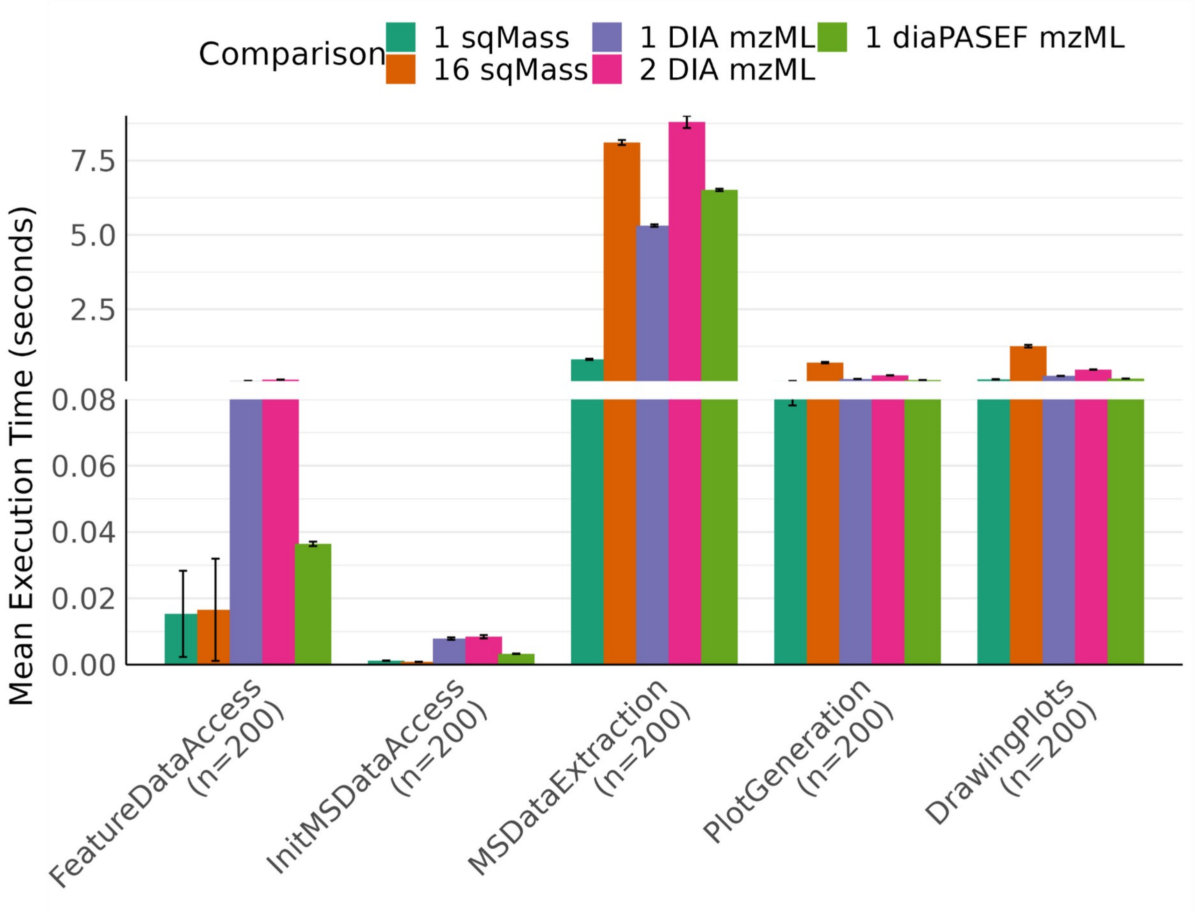

**Supplemental Figure 1. Mean execution time of main processing in MassDash.** Bar plot showing the average execution time with error bars for major processing steps for five different comparison tasks: 1) processing a single post-extracted sqMass run, 2) processing 16 post-extracted sqMass runs, 3) processing a single raw DIA mzML run for targeted extraction, 4) processing two raw DIA mzML runs for targeted extraction, and 5) processing a single raw diaPASEF mzML run for targeted extraction. Data was acquired for each extraction by 200 automated random interactions (precursor selection for extraction) of MassDash. Note: There is a y-axis break to make the lower execution times more visible. Note: Outlier execution time points during the initial load were excluded from the plot to prevent the introduction of extremely large error bars. For detailed information, please refer to supplemental Table 1, where the maximum execution time corresponds to the initial loading of data. Subsequently, the data is cached for expedited access.

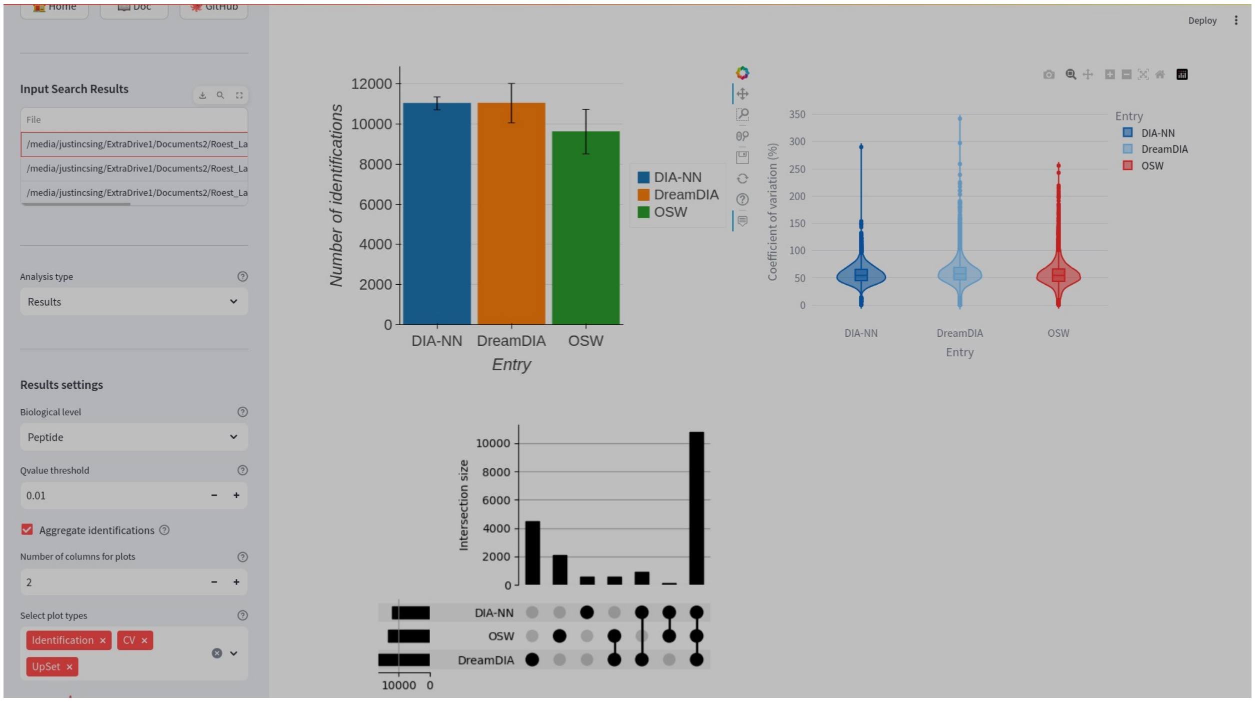

**Supplemental Figure 2. MassDash's Web Interface for Search Results Comparison Analysis.** Search result comparison tab allowing users to input multiple search result files from mass spectrometry data processing software (i.e. OpenSwathWorkflow, DIA-NN, DreamDIA). The example demonstrates a comparison between OpenSwath, DIA-NN, and DreamDIA on the same dataset, displaying an identifications bar plot, a violin plot of coefficient of variation, and an UpSet plot of the intersection comparison in identifications. These resulting plots can be adjusted for biological context (peptide, protein) and FDR threshold. The data used in this example is sourced from the gold standard *S. pyogenes* data, available on PeptideAtlas (PASS01508).

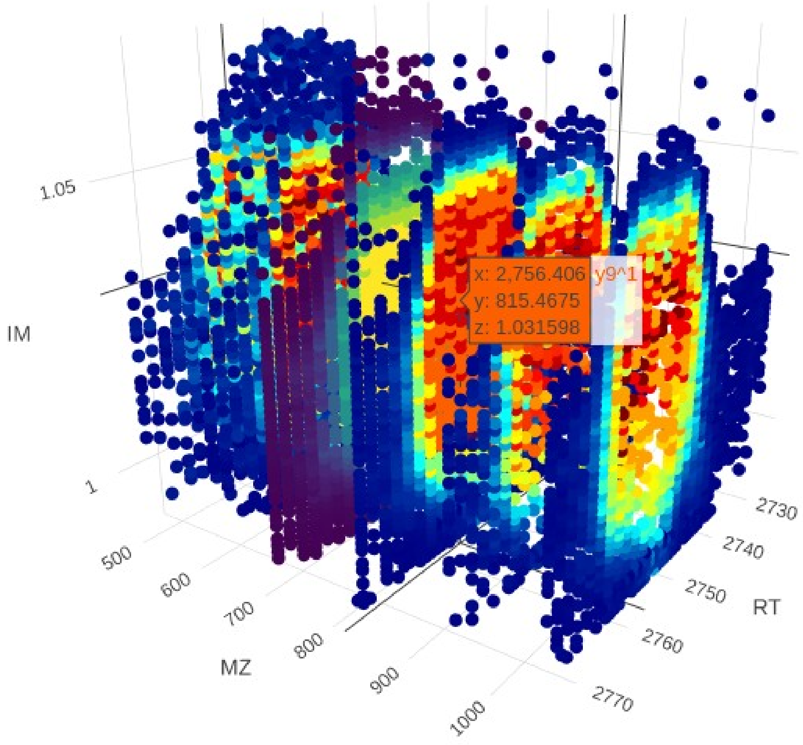

**Supplemental Figure 3. 3D Scatterplot Across Ion mobility, Retention time, and m/z for Precursor *GALQNIIPASTGAAK, charge +2*.** Screenshot of an interactive 3D scatterplot created by MassDash. This example shows that IM heatmaps are highly concordant across the precursor and fragment ions. Hovering over the graph shows the specific point's fragment ion, retention time, m/z, and ion mobility. The plots of the alternative color scheme represent the precursor signal.

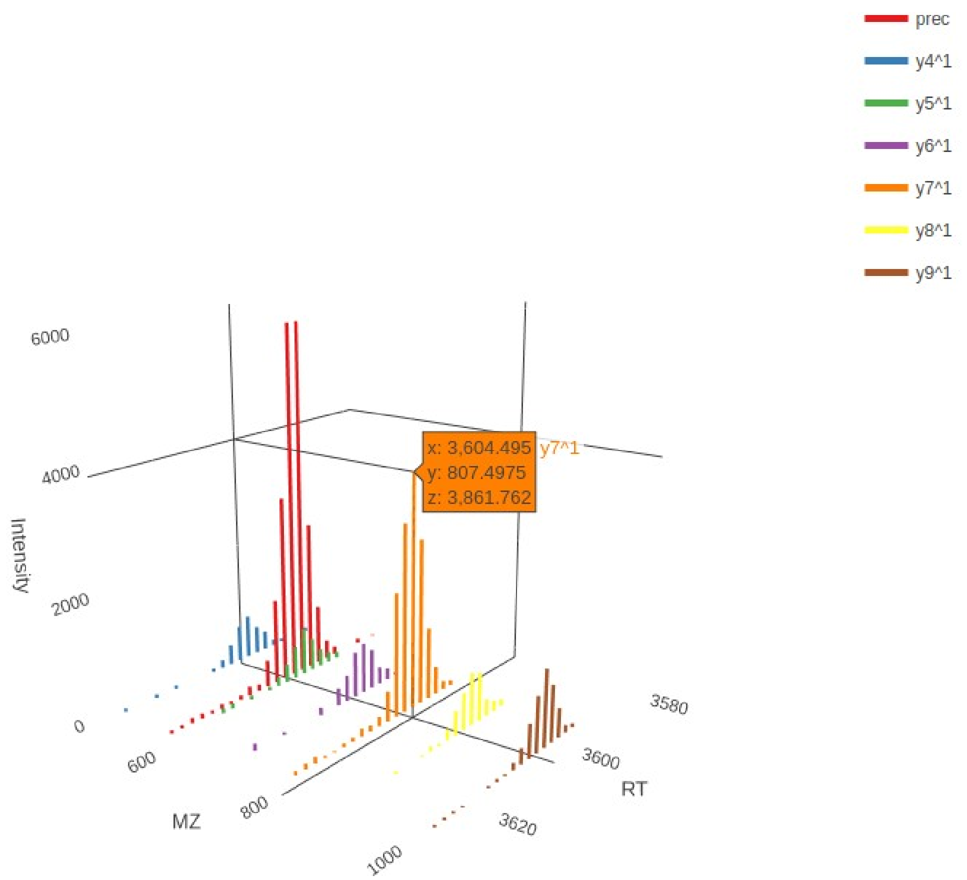

**Supplemental Figure 4. 3D LinePlot Of Precursor *VANVSLLALYK, charge +2*.** Lineplot of all fragments by their m/z. Hovering over the lines records the retention time, m/z, and intensity respectively of the line.

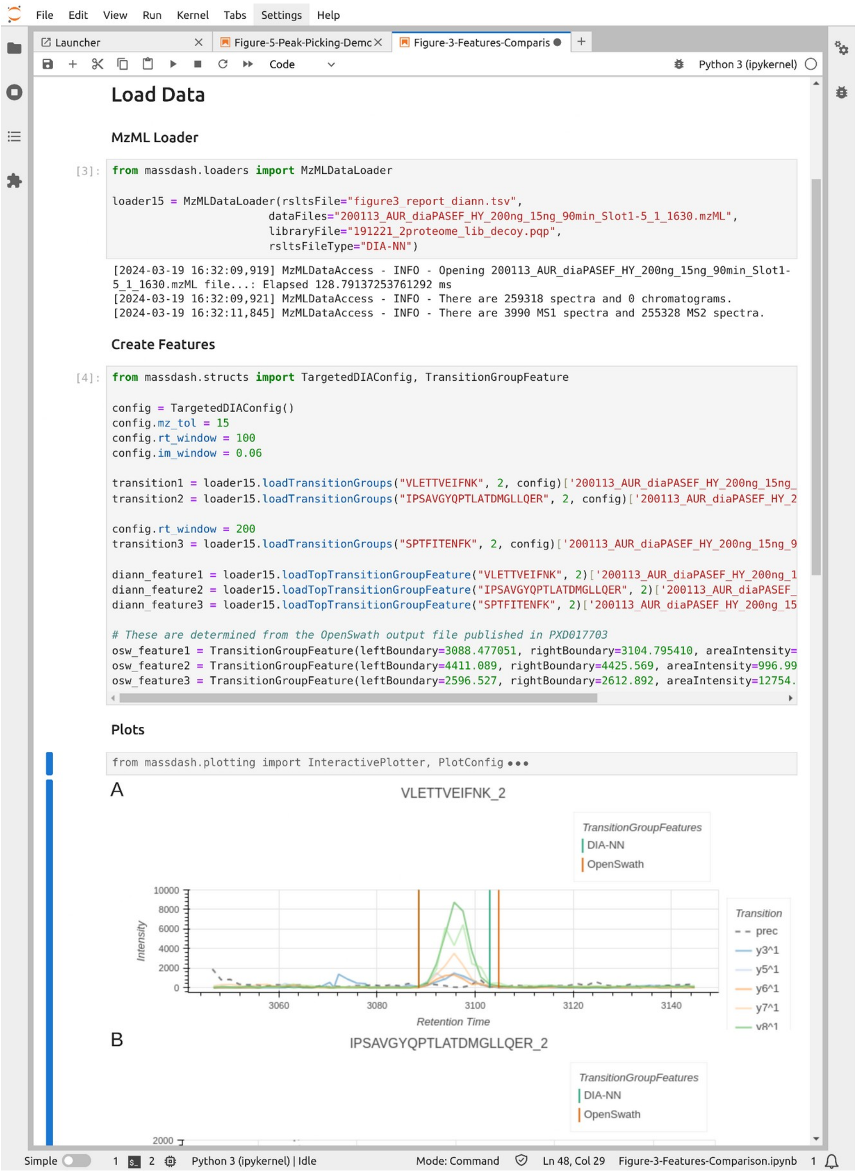

**Supplemental Figure 5. Example of Jupyter Lab Scripting Example.** Screenshot of massDash used in the Jupyter lab context. This specific screenshot is used for plotting Figure 3 of the main text. Note plotting code is hidden.

**Supplemental Table 1. Performance Summary of Memory Usage of MassDash.**

| Memory Performance Summary | | | |
|---|---|---|---|
| **Input Data** | **File Size (Gb)** | **Memory (Gb)[1]** | |
| | | **10 Interactions** | **200 Interactions** |
| 1 sqMass | 0.28 | 2.02 | 2.10 |
| 16 sqMass | 4.4 | 2.12 | 2.59 |
| 1 DIA mzML | 2.4 | 2.57 | 2.45 |
| 2 DIA mzML | 5.7 | 2.70 | 2.62 |
| 1 diaPASEF mzML | 6.1 | 1.83 | 1.88 |

[1] Memory usage is approximated using the maximum amount of resident memory output from `/usr/bin/time` for the entire process of running MassDash GUI with either 10 or 200 random precursor selection for visualization.

**Supplemental Table 2. Performance Summary of Execution Time for Main Processing Steps in MassDash.**

| Performance Summary<br>Summary of performance metrics for each metric | | | |
|---|---|---|---|
| | Execution Time (sec)[1] | | |
| Input Data | Median | Min | Max |
| FeatureDataAccess[2] | | | |
| 1 sqMass | 0.0018 | 0.0004 | 2.5978 |
| 16 sqMass | 0.0009 | 0.0003 | 3.1015 |
| 1 DIA mzML | 0.0867 | 0.0408 | 0.1654 |
| 2 DIA mzML | 0.1287 | 0.0040 | 0.2295 |
| 1 diaPASEF mzML | 0.0335 | 0.0202 | 0.1005 |
| InitMSDataAccess[2] | | | |
| 1 sqMass | 0.0009 | 0.0003 | 3.1414 |
| 16 sqMass | 0.0007 | 0.0003 | 3.2575 |
| 1 DIA mzML | 0.0064 | 0.0037 | 130.9682 |
| 2 DIA mzML | 0.0066 | 0.0022 | 157.3725 |
| 1 diaPASEF mzML | 0.0030 | 0.0017 | 454.0578 |
| MSDataExtraction[2] | | | |
| 1 sqMass | 0.7070 | 0.4668 | 1.7900 |
| 16 sqMass | 7.9660 | 6.3010 | 16.8591 |
| 1 DIA mzML | 5.2633 | 2.5976 | 7.0235 |
| 2 DIA mzML | 9.4220 | 0.0030 | 14.4276 |
| 1 diaPASEF mzML | 6.5129 | 4.5205 | 8.3903 |
| PlotGeneration[2] | | | |
| 1 sqMass | 0.0759 | 0.0051 | 0.2465 |
| 16 sqMass | 0.7614 | 0.0304 | 1.8795 |
| 1 DIA mzML | 0.1491 | 0.0546 | 0.3010 |
| 2 DIA mzML | 0.2666 | 0.1176 | 0.4900 |
| 1 diaPASEF mzML | 0.1121 | 0.0505 | 0.2238 |
| DrawingPlots[2] | | | |
| 1 sqMass | 0.1231 | 0.0019 | 0.4593 |
| 16 sqMass | 1.4173 | 0.0009 | 4.5973 |
| 1 DIA mzML | 0.2500 | 0.1535 | 0.5017 |
| 2 DIA mzML | 0.4614 | 0.2263 | 0.8132 |
| 1 diaPASEF mzML | 0.1564 | 0.0758 | 0.3448 |

[1] Time (execution time) is measured in seconds using wall-clock time of a block of code.

[2] Metrics are calculated from 200 random interactions of MassDash extracting and plotting data for the random precursor.

#### IV. User Manual

##### A. Install

###### The Python Package Index

MassDash is available as a python package on the python package index, and can be installed by `pip` in a terminal or cmd prompt.

```
$ pip install massdash
```

###### Building from Source

The source code is freely open and accessible on Github at https://github.com/Roestlab/massdash. The package can be installed by cloning and installing from source using `pip`.

1. First clone the repository:

```
$ git clone git@github.com:Roestlab/massdash.git
```

2. Change into the massdash directory

```
$ cd massdash
```

3. Install using pip

```
$ pip install -e .
```

###### Docker Image

MassDash is available as a docker image on dockerhub, ensure you have docker installed on your system, you can then pull the latest image of MassDash.

```
$ docker pull singjust/massdash:latest
```

##### B. Usage

MassDash is a modular and flexible python package that has a streamlit graphical user interface (GUI), but can also be used as a python module for regular python scripting. This tutorial will go through the basics of how to use the GUI, for scripting with MassDash, please refer to the API documentation for MassDash.

###### 1. Getting Started

To run the GUI, enter the following in your terminal:

```
$ massdash gui
```

This will redirect you to a browser window under your localhost:port#, the post# by default is usually 8501. If a browser window does not directly open, you can open up a browser window, and enter [http://localhost:8501/](http://localhost:8501/) in the URL search bar (replace 8501 with a different port number reported in the terminal if 8501 is already in use).

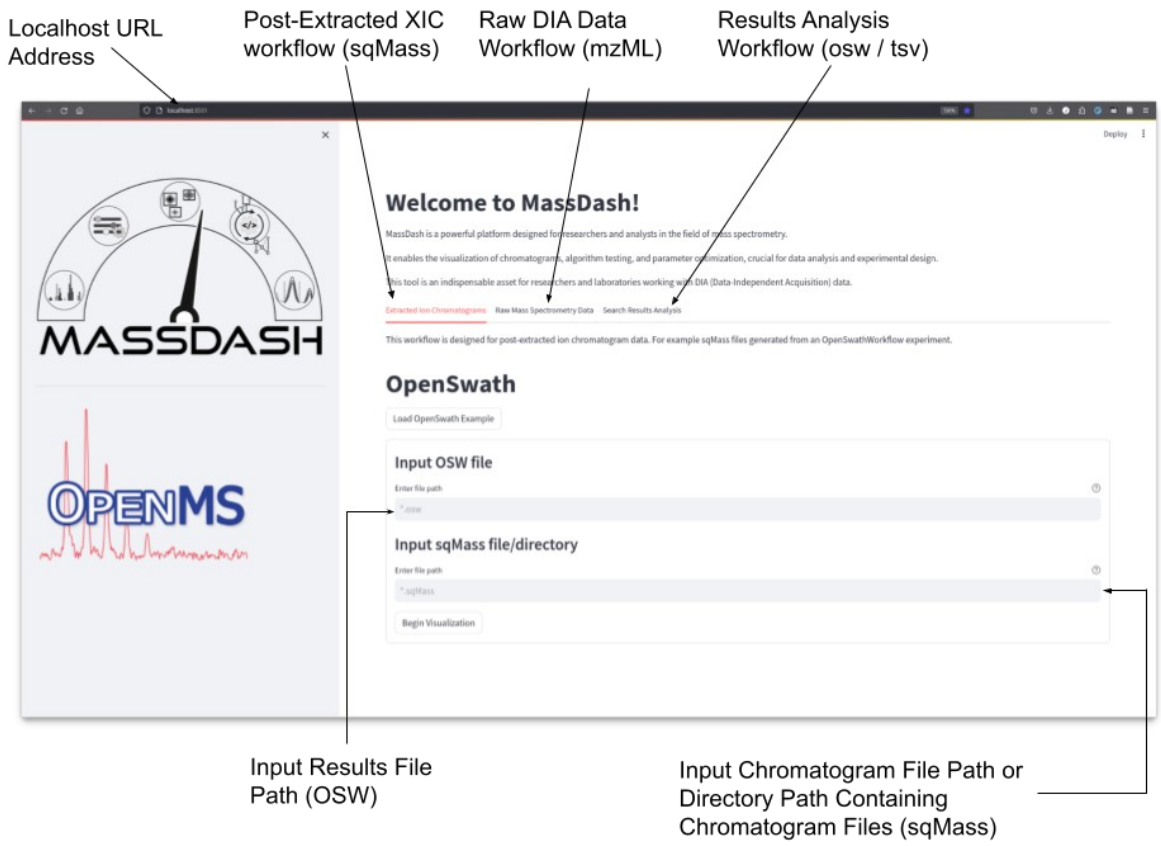

**Figure 1. MassDash GUI Welcome Page.** i) Localhost URL address hosting GUI, ii) Post-extracted XIC workflow tab, iii) Raw DIA data targeted extraction workflow, Iv) Search results analysis workflow tab, v) Results file input for post-extracted XIC workflow, vi) Input XIC sqMass files for post-extracted XIC workflow.

Once you are redirected to the browser url hosting MassDash, you will see a welcome page (Figure 1), that contains three tabs: **1) *Extracted Ion Chromatograms*, 2) *Raw Mass Spectrometry Data*, 3) *Search Results Analysis***. These three tabs are for different workflows, **(1)** allows the user to visualize the resulting extracted ion chromatograms and the peak boundaries identified by an OpenSwathWorkflow analysis, **(2)** allows the user to analyze raw DIA mass spectrometry data on-the-fly, extracting features based on a search results experiment (OpenSwath, DIA-NN, DreamDIA, etc.), and **(3)** allows the user to compare the results of different search result experiments (i.e. comparing different software or comparing different extraction parameters). Each workflow has a *load example data button* that will load test data files, which are available as part of the github repository, or alternatively, you can enter the file paths for the input data, and press the *Begin Visualization* button to load the data and workflow.

#### 2. Post-Extracted Ion Chromatogram Workflow

Upon entering the input files and starting the workflow **(1)**, you will see the sidebar populated with several elements including: **1.1) input file information, 1.2) transition list information, 1.3) plot control settings, and 1.4) peak picking settings**. The main area will be populated with interactive Bokeh figures and a drop-down logging text area that display time-execution information on different processes that were executed for extracting, generating and rendering the extracted ion chromatogram (XIC) figures (Figure 2).

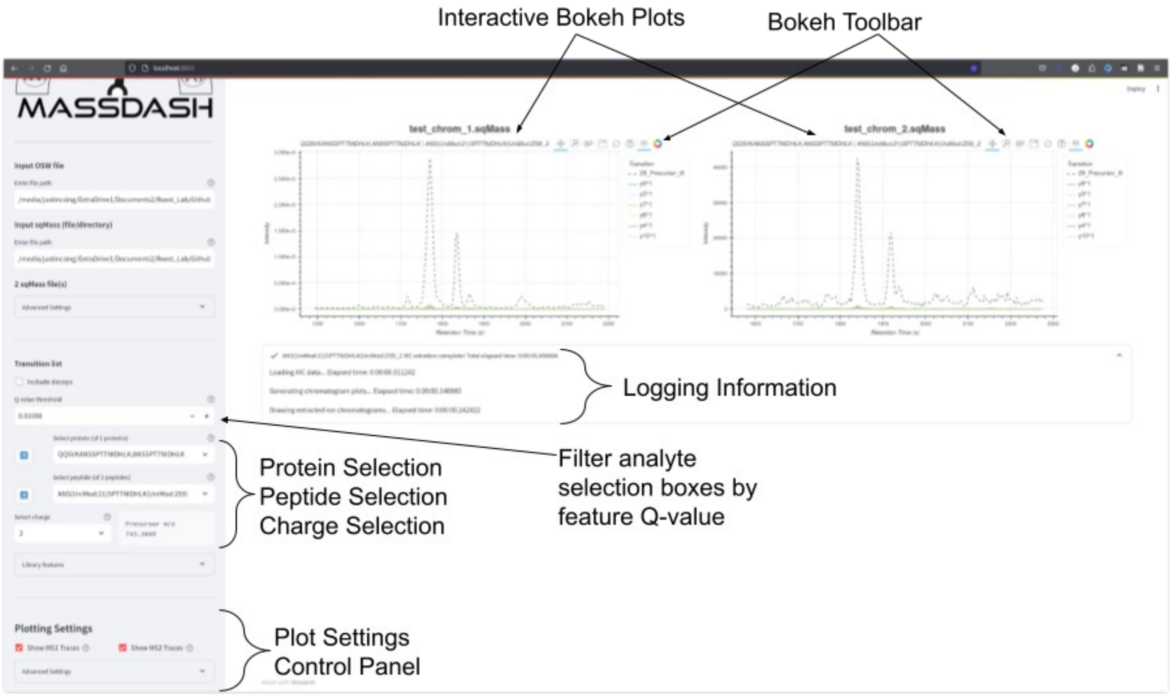

**Figure 2. Extracted Ion Chromatograms Workflow Example.** Example showcasing extracted ion chromatograms for two runs. The main area contains interactive Bokeh plots, and the sidebar contains analyte selection settings and plot control settings.

Each figure contains a toolbar that allows for interactive panning, zooming, information hovering, and figure saving. In addition, each plot has an interactive legend that allows for muting of individual traces. The sidebar contains several sections that allow for controlling for analyte selection and control for plotting attributes. The analyte selection **(1.2)** allows for selecting which analyte to plot, and is split into individual protein, peptide and charge searchable drop down selection boxes. You can either select the analyte via the selection dropdown or you can use the random selection buttons that will randomly select an analyte to plot. By default, the analytes populated in the drop down selection boxes are filtered based on the feature Q-value of 1%, this can be adjusted if you want to include less confident or more confident identifications.

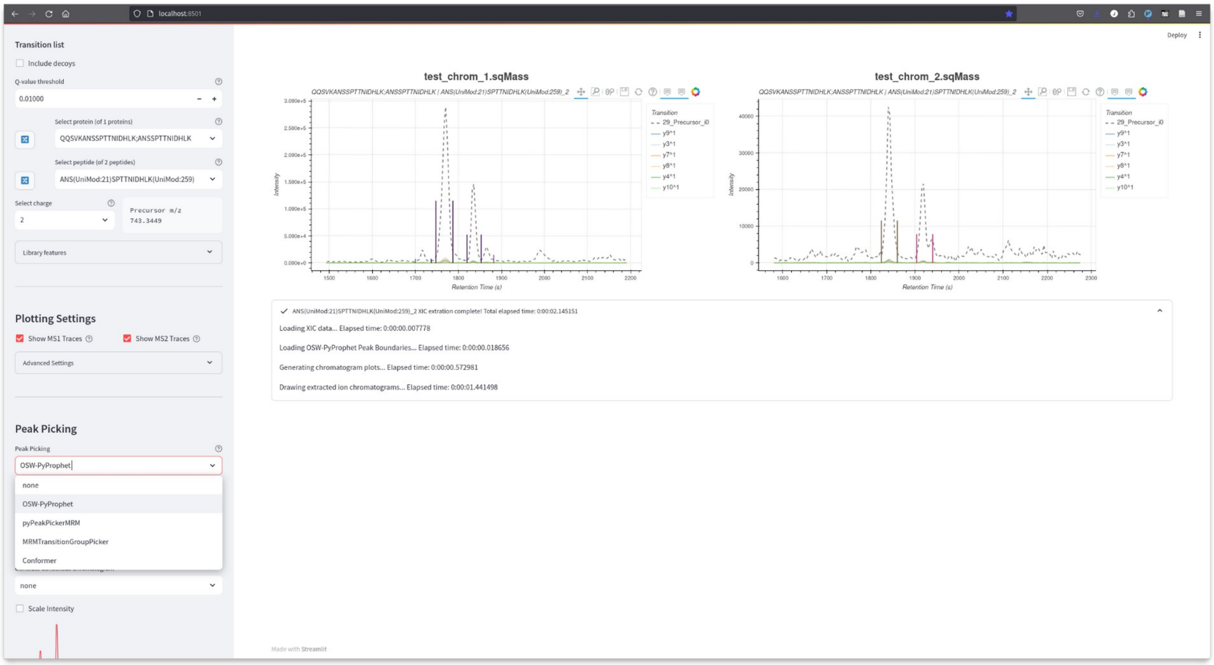

**Figure 3. Peak-Picking Dropdown List Options.** The peak-picking dropdown selection allows the user to select different peak-picking algorithms to display peak boundaries on the chromatogram figure.

The figures can be further controlled, by adjusting settings in the Plotting Settings section, that allows for displaying or hiding MS1 or MS2 traces. The advanced dropdown panel allows for controlling how the plots are arranged in a grid and allows for smoothing of the traces. Aside from just visualizing the XICs, it is also possible to visualize the peak boundaries identified by OpenSwath. This can be turned on in the Peak-Picking dropdown selection box, which allows for the selection of either the identified peaks by OpenSwath or on-the-fly peak picking with several different peak-picking algorithms (Figure 3). Once selected, the peak boundaries will be rendered onto the figure. Each peak boundary contains meta-information that can be hovered over to show information about the identification, such as the peak boundaries, the retention time apex, the retention time apex intensity, and the peak group features Q-Value (Figure 4).

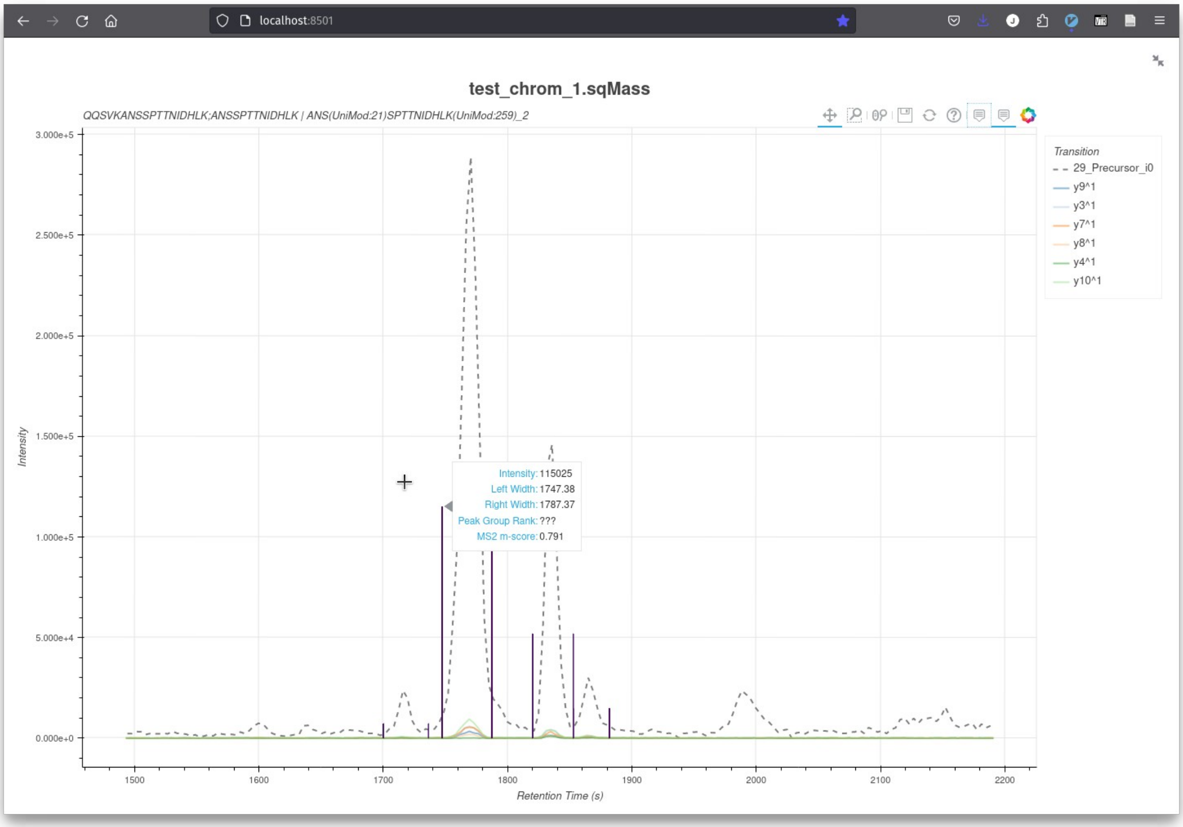

**Figure 4. Peak Boundary Information.** Hovering over the peak boundaries displays text information about the identified peak.

#### 3. On-The-Fly Targeted Extraction of Raw DIA Mass Spectrometry Data Workflow

Once you select the ***Raw Mass Spectrometry Data*** tab, you will see the workflow depicted in Figure 5 illustrating the input components for the Raw Targeted Data Extraction process. These inputs include: i) the file path of the transition list containing analyte information, ii) the file or directory path containing raw Data-Independent Acquisition (DIA) mass spectrometry data, iii) the file path for search result files containing feature identification results, and iv) a dropdown selection to specify the software from which the search results were derived. These elements collectively form the essential parameters for executing targeted data extraction, providing a comprehensive view of the key inputs guiding the workflow.

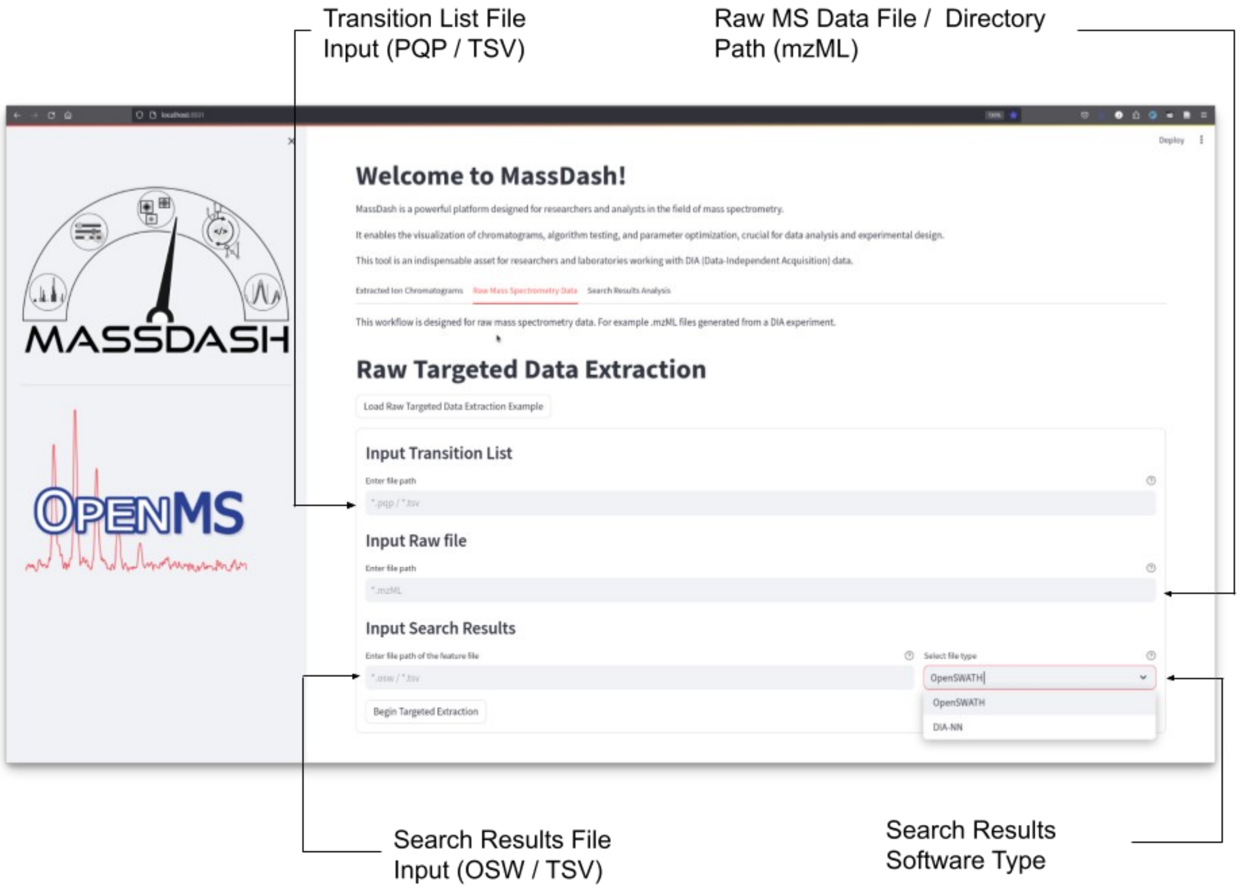

**Figure 5. Raw Targeted Data Extraction Workflow Input.** i) Transition list file path containing analyte information, ii) file or directory path containing raw DIA mass spectrometry data, iii) file path for search result files containing feature identification results, and iv) dropdown selection to choose what software the search results were derived from.

In Figure 6, a practical illustration of targeted extraction for diaPASEF data is presented. The sidebar panel showcases controls for selecting analytes from the transition list, a dropdown text area with search result details derived from DIA-NN for the chosen precursor. The main panel provides visualizations of the extraction ion chromatogram and the extracted ion mobilogram, utilizing identified retention time and ion mobility coordinates from DIA-NN. The accompanying table below the plots contains raw extracted data, offering the option to save the information as a CSV file for subsequent analysis and manipulation. The extraction parameters in the sidebar allows the user to adjust the mass-to-charge tolerance (ppm) window, the retention time window and the ion mobility window settings for optimizing the extraction (Figure 7).

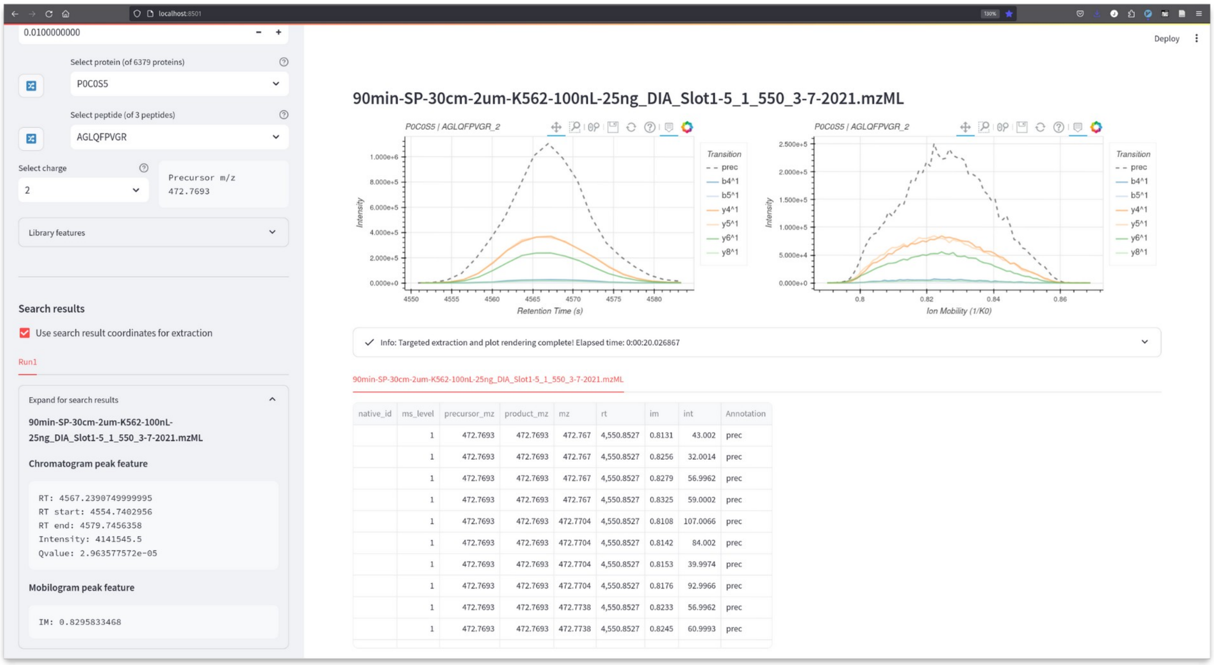

| native_id | ms_level | precursor_mz | product_mz | mz | rt | im | int | Annotation |
|---|---|---|---|---|---|---|---|---|
| | 1 | 472.7693 | 472.7693 | 472.767 | 4,550.8527 | 0.8131 | 43.002 | prec |
| | 1 | 472.7693 | 472.7693 | 472.767 | 4,550.8527 | 0.8256 | 32.0014 | prec |
| | 1 | 472.7693 | 472.7693 | 472.767 | 4,550.8527 | 0.8279 | 56.9962 | prec |
| | 1 | 472.7693 | 472.7693 | 472.767 | 4,550.8527 | 0.8325 | 59.0002 | prec |
| | 1 | 472.7693 | 472.7693 | 472.7704 | 4,550.8527 | 0.8108 | 107.0066 | prec |
| | 1 | 472.7693 | 472.7693 | 472.7704 | 4,550.8527 | 0.8142 | 84.002 | prec |
| | 1 | 472.7693 | 472.7693 | 472.7704 | 4,550.8527 | 0.8153 | 39.9974 | prec |
| | 1 | 472.7693 | 472.7693 | 472.7704 | 4,550.8527 | 0.8176 | 92.9966 | prec |
| | 1 | 472.7693 | 472.7693 | 472.7738 | 4,550.8527 | 0.8233 | 56.9962 | prec |
| | 1 | 472.7693 | 472.7693 | 472.7738 | 4,550.8527 | 0.8245 | 60.9993 | prec |

**Figure 6. Example Targeted Extraction for diaPASEF data.** Sidebar panel demonstrates transition list controls for analytes selection, and search results information obtained from DIA-NN for the selected precursor. The main panel displays the extraction ion chromatogram and the extracted ion mobilogram for the current precursor based on identified retention time and ion mobility coordinates from DIA-NN. The table below the plots is the raw extracted data which can be saved to a csv file for further manipulation.

The benefit of extracting from the raw data is the direct access to the full extracted data for a selected analyte. This allows the user to plot different kinds of plots and different dimensions of plots, one dimension plots, two dimension plots and three dimension plots (Figure 7). For one dimensional plots, the user can visualize the extracted spectra, chromatogram and mobilogram. Peak-picking can also be applied as in the first workflow, specifically for the extracted ion chromatograms. Two dimensional plots allow heatmap style visualizations of two dimensions, i.e. retention time vs ion mobility.  Three dimensional plots allow you to visualize spectrum-chromatogram plots, scatter heatmap plots of mass-to-charge (Figure 7), retention time and ion mobility, and three dimensional contour plots of two dimensions.

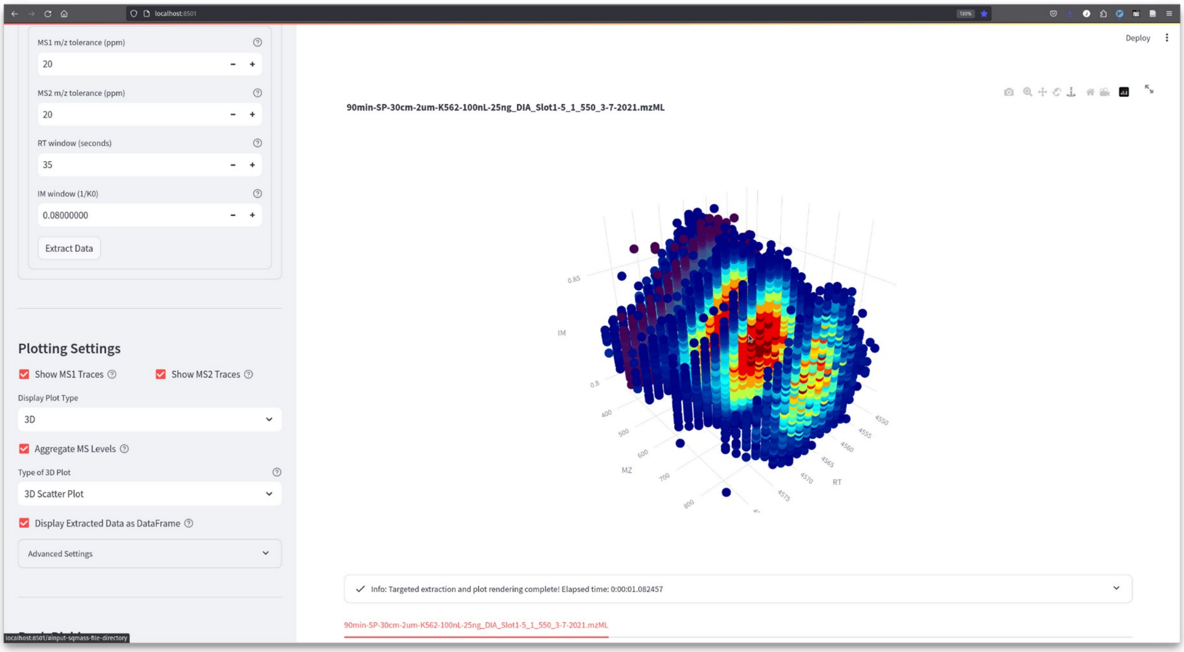

**Figure 7. Extraction Parameters and Different Types of Plots.** The dropdown extraction parameters allows for optimizing the extraction windows for mass-to-charge, retention time and ion mobility. The plot settings in the sidebar panel allows for selecting between different types of plots, one dimension plots, two dimension plots and three dimension plots.

#### 4. Search Results Comparisons Analysis Workflow

Once the ***Search Results Analysis*** tab is selected, you will see two text input widget fields, and a dropdown selection bow (Figure 8). The text input fields are for the search results file path, a label to name the experiment for the search results, and the dropdown selection box is used to determine what software generated the search results. Additional search results can be added by toggling the numeric input widget to add additional entries. There are two different types of analysis workflows currently supported, a results workflow and a score distributions workflow, which is selected in the dropdown box in the sidebar. The default workflow is set to the results workflow.

In Figure 9, the analysis of search results comparisons is depicted, highlighting the versatility of MassDash. The sidebar provides settings to control results at a specified Q-value cutoff, enabling the generation of informative visualizations. These include an identifications bar plot, a log2 quantifications violin plot, a coefficient of variation violin plot, and an upset comparisons plot. This figure underscores MassDash's capability to facilitate comprehensive and customizable comparisons at the biological level, empowering researchers in their data interpretation and analysis. The analysis of score distributions provides users with a visual representation of the distributions of targets and decoys across various scoring levels and score variables. The main panel displays Bokeh plots illustrating the score distributions, accompanied by a table presenting the corresponding feature scores and variables (Figure 10).

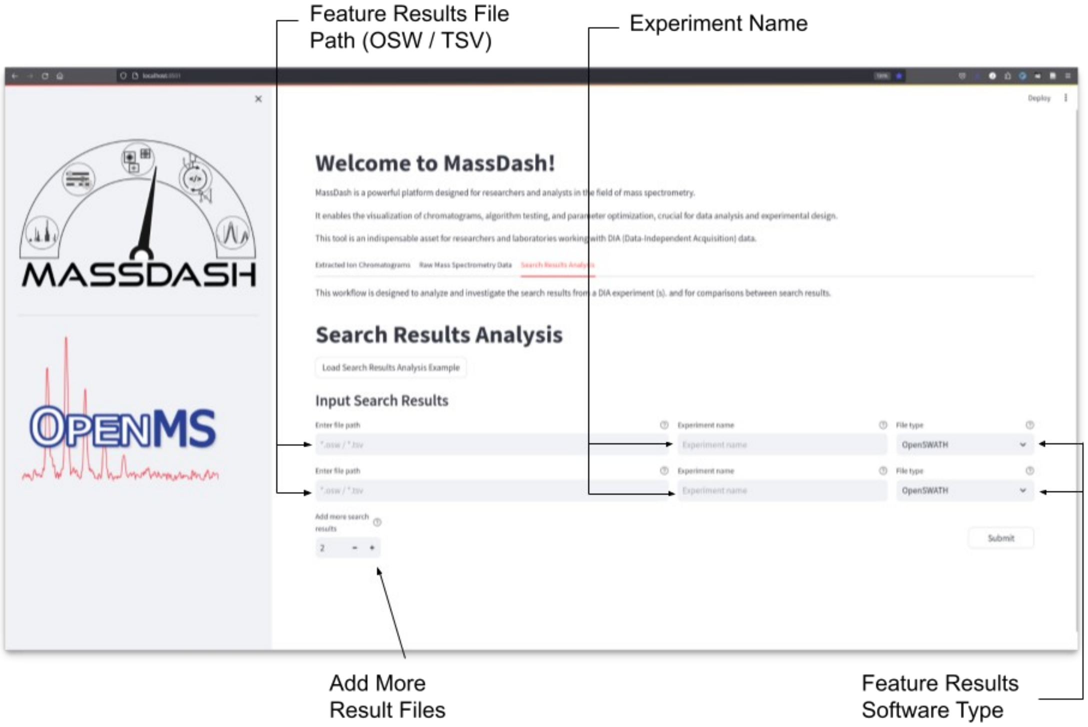

**Figure 8. Search Results Analysis Comparison Workflow Input.** The welcome page for the search results analysis tab required a feature results file path, an experiment label, and the type of software from the dropdown that generated the results. Additional search results can be added from the numeric input toggle button.

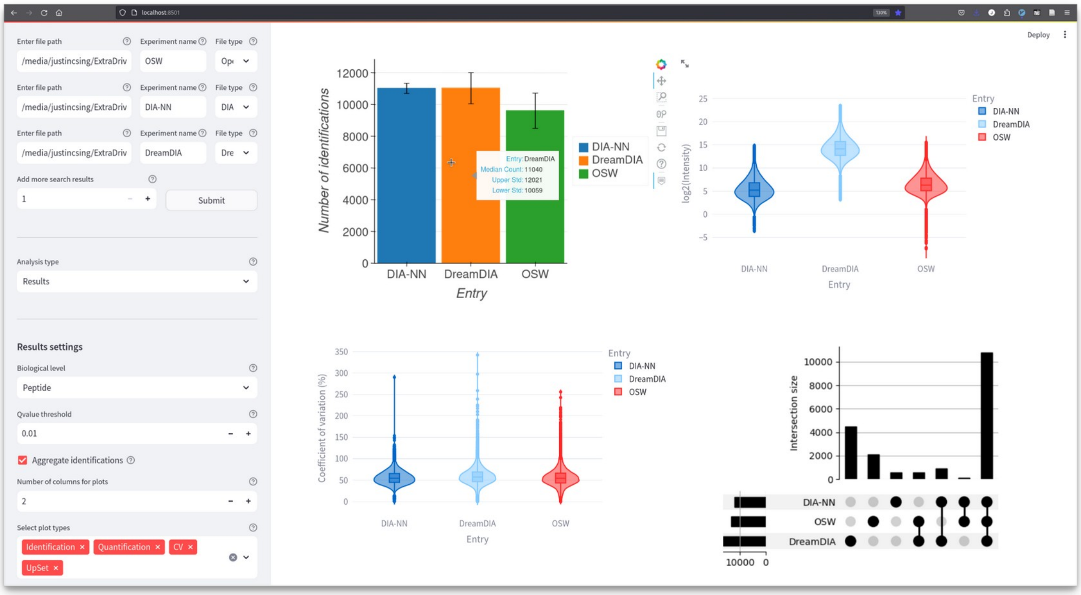

**Figure 9. Search Results Comparisons Analysis.** Sidebar results settings allow for control on the biological level at a specific Qvalue cutoff to generate an identifications bar plot, a log2 quantifications violin plot, a coefficient of variation violin plot, and an upset comparisons plot.

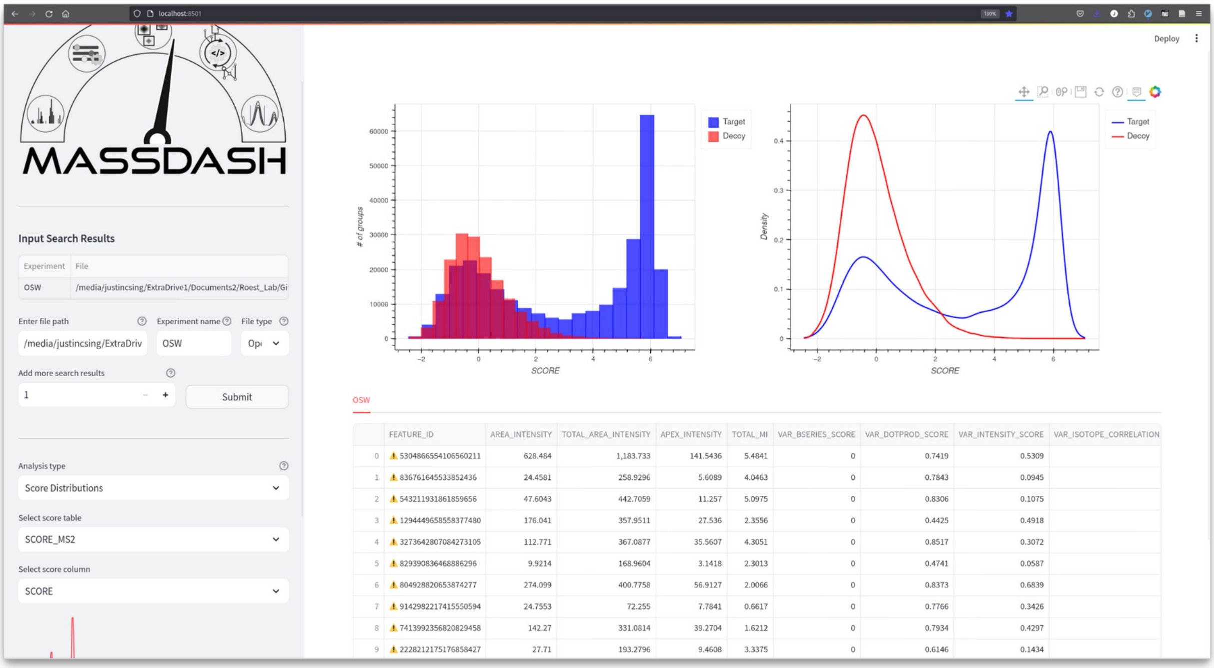

**Figure 10. Search Results Score Distributions.** The score distributions analysis allows the user to visualize the distributions of targets vs decoys for different scoring levels and different score variables. The main area displays bokeh plots of the distribution of scores and a table of the feature scores.
